# Gentrification influences mosquito community composition at neighborhood and county levels in Miami-Dade County, Florida

**DOI:** 10.1101/2025.04.30.651397

**Authors:** Nicole A. Scavo, Chalmers Vasquez, Laura C. Multini, John-Paul Mutebi, André B. B. Wilke

## Abstract

Gentrification is occurring across urban areas in the United States and poses threats to marginalized and vulnerable communities through displacement, disruption of social networks, and worsening health outcomes. Gentrification is both a social and environmental process, affecting socioecological factors responsible for driving mosquito abundance and community composition. Our study aims to investigate how gentrification in Miami-Dade County, Florida, affects the alpha and beta diversity of mosquito communities. We relied on data from the Miami-Dade County Mosquito Control Division from 2020 to 2024, paired with data from the American Community Survey, to analyze changes in mosquito community composition based on gentrification status. Our results show that gentrification, measured by changes in home value, age, race, and education, significantly affected mosquito richness and community composition at county and neighborhood levels. *Culex quinquefasciatus* and *Aedes aegypti*, primary arbovirus vector species, were more abundant in gentrifying areas, representing 31% of community composition variation compared to non-gentrifying areas. These findings have important implications for improving mosquito-borne disease preparedness and response in urban settings.

## Introduction

Over half of the global population lives in urban areas (1). Urban areas are predicted to expand and occupy more land throughout this century (2) with socioecological factors playing an important role in how this expansion unfolds (3). Gentrification (i.e., economic and demographic change in historically disinvested neighborhoods) is an example of the intersection of socioecological processes shaping human communities, public health, and diversity within urban areas. Gentrification disproportionately affects vulnerable and underserved communities, particularly communities of color (4,5). Between 2000 and 2013, approximately 110,000 Black and 24,000 Hispanic families were displaced in the U.S. due to gentrification (6). Gentrification influences health outcomes by altering access to affordable housing, healthy food, social networks, and exercise facilities, often leading to displacement and associated health risks (7–11). It has also been linked to changes in disease incidence in urban areas due to shifts in human behavior, movement, and increased green space and habitat connectivity (12,13). Recognizing these impacts, the U.S. Centers for Disease Control and Prevention designated gentrification as a public health issue in 2009 (14).

Mosquito community composition, presence, and abundance are influenced by socioecological factors (15–19). Species-specific ecological and physiological traits determine how mosquitoes respond to habitat variations and resource availability variations (18,20–23). The loss of key resources commonly found in natural habitats can lead to mosquito population declines or local extinctions of species unable to rely on resources available in urban environments. In contrast, resource availability in urban environments can increase vector species abundance, potentially increasing the risk of pathogen transmission (24). Moreover, gentrification affects mammal diversity (25), potentially altering host availability for mosquito vectors, increasing mosquito-human contact, and the risk of pathogen spillover (26).

Miami-Dade County, Florida, is a major arbovirus gateway for arbovirus entry to the United States (27,28). In 2016, public health authorities responded to a local Zika virus outbreak, and between 2010 and 2024, 314 cases of locally transmitted dengue were reported in the county (29). However, according to CDC estimates, a multiplication factor of 21-105 is needed to correct for the under-reporting of the number of laboratory-positive dengue inpatients (i.e., reporting rate of 1.0%-4.8%) (30). This suggests that dengue virus transmission within the United States is far higher than what is reported in the official records. Miami-Dade County has a diverse population, including minority and underserved groups. Current predictions indicate that approximately 700,000 people are expected to move to Miami-Dade County by 2030, further exacerbating gentrification. Furthermore, climate gentrification is predicted to displace over 50% of residents if sea levels rise less than 1 meter (31).

Although gentrification is a significant anthropogenic driver of environmental change, its impact on the community composition and abundance of mosquito vector species in Miami-Dade County, Florida, remains unknown. We hypothesize that gentrified areas will have lower species richness, whereas suburban and rural areas will support greater mosquito diversity. Additionally, we hypothesize that mosquito community composition will differ based on gentrification status, with these variations primarily driven by two vector species, *Culex quinquefasciatus* and *Aedes aegypti* (32). Therefore, our objective was to investigate the effects of gentrification on mosquito presence, abundance, and community composition to better understand its impact on the risk of arbovirus transmission in urban areas.

## Methods

### Site selection

Miami-Dade County, Florida, is home to a population of approximately 2.6 million people (33) and covers an area of about 6,000 km^2^. Miami-Dade County has a tropical climate with average temperatures ranging from an average high of 91°F in summer months and an average low of 61°F in the winter months and average monthly precipitation ranging from 1.8 to 10.5 inches (34). Gentrification has been occurring as an ongoing issue in Miami-Dade County (35,36). *Aedes aegypti* and *Cx. quinquefasciatus* are abundant year-round in Miami-Dade County and local transmission of dengue, Zika, and West Nile viruses has occurred in the county within the past 10 years (27,29,37).

### Mosquito collection

The Miami-Dade Mosquito Control Division’s surveillance system was designed to have at least one mosquito trap per 1.6 km^2^ in urbanized areas, with additional traps around the city limits and points of interest where residents or tourists are likely to be exposed to mosquito vectors. During the period of our study, from January 2020 to December 2024, 320 mosquito traps (282 BG traps and 38 CDC light traps) were set weekly and baited with CO_2_ produced by 1 kg of dry ice pellets (38). All collected mosquitoes were morphologically identified at the Miami-Dade County Mosquito Control Laboratory using morphological keys (39). Male mosquitoes are not attracted to CO_2_-based traps and were considered accidental catches and excluded from analyses.

### Gentrification Variables & Neighborhood Selection

We extracted sociodemographic variables from the American Community Survey and ESRI (40,41) at a 0.5 km buffer radius around each trap location. The following variables were used to create a gentrification index: median home values (2022, 2024), median household income (2022, 2024), median age (2020, 2024), median educational attainment base (2022, 2024), population density (2020, 2024), and race data (Hispanic, Not Hispanic, White, Black, Asian, Native Hawaiian, Other, two or more races in 2022 and 2024). Years were chosen to match the years of mosquito collection, if 2020 data was not available for a specific category, then 2022 data was used. These variables were chosen due to their common use in other studies on gentrification (42,43).

Changes in these variables were used to indicate gentrification. For instance, an increase in home values and a decrease in minority populations would indicate gentrification. Change variables were scaled and then added together to create a composite gentrification index. This index was used to classify individual trap locations (0.5 k buffer around each trap location) at the county level as gentrifying or not gentrifying, with the top quantile of the index being classified as gentrification (Figure 1). This index was also used as the basis for selecting neighborhoods for the neighborhood-level analysis. Six neighborhoods (3 gentrifying and 3 non-gentrifying) were chosen based on similar characteristics: Wynwood, Kendall, North Miami Beach, South Miami Beach, Civic Center, and Westchester. At least 8 traps from each neighborhood were included in the analyses.

**Figure 1.**
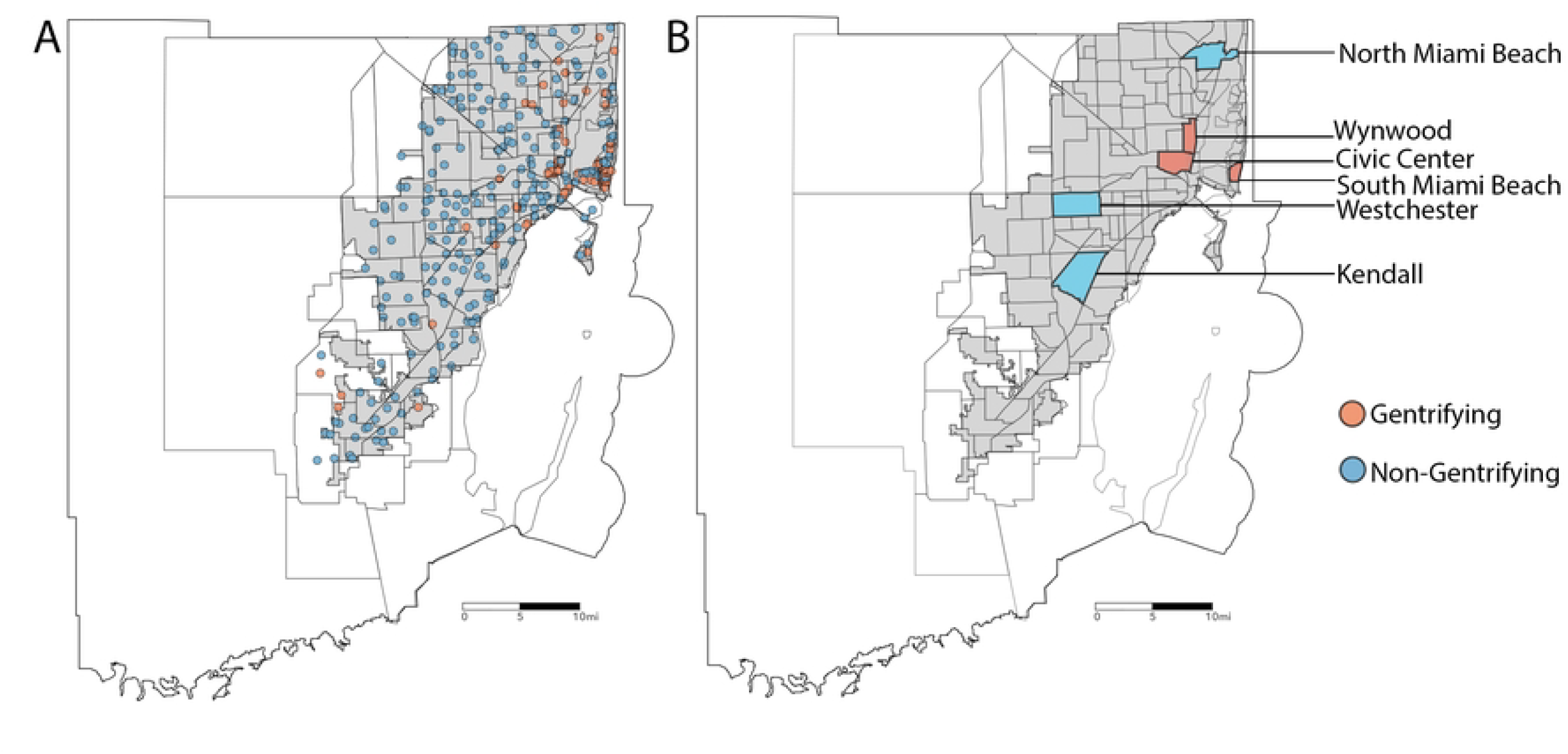
Trap locations and gentrification status in Miami-Dade County. A. Locations of mosquito traps, with blue indicating traps placed in areas below the gentrification threshold and red indicating traps in areas above the threshold. B. Neighborhoods classified by gentrification status, with blue representing non-gentrifying neighborhoods and red representing gentrifying neighborhoods.

### Statistical Analysis

Analyses were conducted at the county and neighborhood levels to identify gentrification variables influencing mosquito richness and to assess differences in mosquito communities between gentrifying and non-gentrifying areas. First, we ran generalized linear mixed models (GLMMs) at the county and neighborhood levels. This approach was chosen due to GLMMs ability to handle unbalanced data sets and to control for temporal factors by using random effects (44,45). Poisson, negative binomial, and quasi-Poisson distributions were tried at each level, and the best-fit distribution was chosen based on minimization of AIC values and assessment of QQ plots. Explanatory variables were scaled before model creation. Backward elimination was then used to choose the best-fit model. For the county-wide analyses, fixed factors included change in home value, change in population density, change in age, change in education, change in the percent of the population that is Black, and change in the percent of the population that is Hispanic. The fixed effects were sampling month (i.e., the cumulative month of the sampling period) to account for differences due to seasonality and trap to account for potential spatial correlation. The fixed effects were the same at the neighborhood level, and the random effects were sampling month and pair identification. The following communities were paired together: Wynwood and Kendall, North Miami Beach and South Miami Beach, and Civic Center and Westchester.

We also compared community composition, or beta diversity, using a permutational multivariate analysis of variance (perMANOVA), a permutational version of the classic MANOVA that tests for the differences in the centroids among groups (46). We used a Bray-Curtis dissimilarity matrix with 10,000 permutations to test the null hypothesis that there were no differences in beta diversity between gentrifying and non-gentrifying traps at the county and neighborhood levels. To complement this analysis, we ran a similarity percentage analysis (SIMPER) to determine which mosquito species were driving the differences in community composition. Lastly, we calculated the Jaccard’s Index as a measure of turnover (47). This was done by comparing community composition from 2020 to 2024 at both the county and neighborhood levels. A Jaccard dissimilatory matrix was used on presence-absence data, and then a mean was calculated and subtracted from one to determine turnover.

## Results

Between 2020-2024, a total of 2,542,150 female mosquitoes were collected in Miami-Dade County. Of these, 1,232,569 were collected using CDC traps and 1,309,581 were collected using BG-Sentinel traps. The six most abundant species were *Cx. quinquefasciatus* (798,018 females), *Culex nigripalpus* (689,848 females), *Aedes taeniorhynchus* (272,358 females), *Ae. aegypti* (269,727 females), *Aedes tortilis* (160,238 females), and *Wyeomyia vanduzeei* (79,211 females).

At the county level, the global GLMM with a negative binomial distribution had the best fit (AIC = 246,214.8). After backward elimination using minimization of AIC, the best-fit model (AIC = 246,214.5) included three explanatory variables (Table 1) and explained 27.9% of the variation in the data when controlling for the effects of the month (i.e., seasonality) and trap location. Overall, traps associated with gentrification variables had lower species richness. Change in home value from 2022 to 2024 had a negative but small effect on mosquito species richness (Exponentiated Estimate: 0.94, 95% Confidence Interval: 0.91 – 0.97, p < 0.001). Population density changes also had a small and negative effect on species richness (Exponentiated Estimate: 0.94, 95% Confidence Interval: 0.90 – 0.97, p < 0.001). Change in the percent Hispanic population had a positive effect on mosquito species richness, with areas with a higher percent Hispanic population being 1.06 times more likely to have higher mosquito richness (Exponentiated Estimate: 1.06, 95% Confidence Interval: 1.02 – 1.09, p = 0.001).

**Table 1.** Best fit generalized linear mixed model (negative binomial distribution) at the county level showing variables contributing to differences in mosquito richness in gentrifying vs non-gentrifying areas. Cumulative month and TrapID were used as random effects. Conditional R^2^ = 0.292. Significant *p* values have been bolded.

| Predictor | Estimate | Exp.<br>Estimate | 95% CI | $p$ |
| --- | --- | --- | --- | --- |
| intercept | 0.71 | 2.04 | 1.91 – 2.18 | **<0.001** |
| Scaled change in home value | -0.06 | 0.94 | 0.91 – 0.97 | **<0.001** |
| Scaled change in population density | -0.06 | 0.94 | 0.90 – 0.98 | **0.002** |
| Scaled change in percent of population that is Hispanic | 0.05 | 1.06 | 1.02 – 1.09 | **0.001** |

The perMANOVA showed that community composition differed at the county level based on gentrification (F = 3.92, p = 0.02; Table 2). These differences were driven by *Cx. quinquefasciatus* and *Ae. aegypti*, which explained 23% and 8% of the differences between communities, respectively (Table 2). Both species were more abundant in gentrifying areas. Lastly, species turnover was similar between gentrifying and non-gentrifying areas, with both yielding a value of 0.43 (Table 2).

**Table 2.** Results for community composition analyses: perMANOVA, SIMPER, and turnover index at the county level. Significant *p* values have been bolded.

| perMANOVA |  |  |  |  |  |  |  |
| --- | --- | --- | --- | --- | --- | --- | --- |
|  | df | SumOfSqs | R<sup>2</sup> | F | p |  |  |
| Model | 1 | 0.70 | 0.01 | 3.92 | 0.02 |  |  |
| Residual | 298 | 53.11 | 0.99 |  |  |  |  |
| SIMPER |  |  |  |  |  |  |  |
| Species | average | sd | ratio | ava | avb | cumsum | p |
| *Cx. quinquefasciatus* | 0.23 | 0.18 | 1.25 | 770 | 533 | 0.44 | 0.002 |
| *Ae. aegypti* | 0.08 | 0.07 | 1.08 | 208 | 212 | 0.60 | 0.08 |
| *Cx. nigripalpus* | 0.06 | 0.14 | 0.38 | 244 | 575 | 0.70 | 0.76 |
| *Ae. tortilis* | 0.04 | 0.11 | 0.38 | 38 | 301 | 0.78 | 0.74 |
| *Wy. vanduzeei* | 0.04 | 0.08 | 0.47 | 26 | 76 | 0.5 | 0.94 |
| *Ae. taeniorynchus* | 0.03 | 0.09 | 0.33 | 43 | 242 | 0.91 | 0.99 |
| **Turnover Index** |  |  |  |  |  |  |  |
| Gentrifying | 0.43 |  |  |  |  |  |  |
| Non-gentrifying | 0.43 |  |  |  |  |  |  |

We conducted the same analyses at the neighborhood level. The GLMM showed that mosquito richness was influenced by changes in home value, population density, age, the percent of the population that is Black, and the percent of the population that is Hispanic (Table 3). Increases in home values (Exponentiated Estimate: 0.87, 95% Confidence Interval: 0.85 – 0.88, p < 0.001) and population density (Exponentiated Estimate: 0.93, 95% Confidence Interval: 0.92 – 0.94, p < 0.001) were associated with lower mosquito richness, though the effects were small. A decrease in age was also associated with lower mosquito richness (Exponentiated Estimate: 0.98, 95% Confidence Interval: 0.97 – 0.99, p = 0.002). Decreases in both Black (Exponentiated Estimate: 1.13, 95% Confidence Interval: 1.11– 1.15, p < 0.001) and Hispanic populations (Exponentiated Estimate: 1.22, 95% Confidence Interval:1.20– 1.25, p < 0.001) were associated with lower richness.

**Table 3.** Best fit generalized linear mixed model (Poisson distribution) at the neighborhood level showing variables contributing to differences in mosquito richness in gentrifying vs non-gentrifying areas. Cumulative month and pairID were used as random effects. Conditional R^2^ = 0.17. Significant *p* values have been bolded.

| Predictor | Estimate | Exp.<br>Estimate | 95% CI | $p$ |
| --- | --- | --- | --- | --- |
| intercept | 0.76 | 2.13 | 1.83-2.48 | **<0.001** |
| Scaled change in home value | -0.14 | 0.87 | 0.85 – 0.88 | **<0.001** |
| Scaled change in population density | -0.08 | 0.93 | 0.92 – 0.94 | **<0.001** |
| Scaled change in age | -0.02 | 0.98 | 0.97 - 0.99 | **0.002** |
| Scaled change in percent of population that is Black | 0.12 | 1.13 | 1.11 – 1.15 | **<0.001** |
| Scaled change in percent of population that is Hispanic | 0.20 | 1.22 | 1.20 – 1.25 | **<0.001** |

The community composition analyses at the neighborhood level showed that mosquito communities were different between gentrifying and non-gentrifying neighborhoods (F = 4.7, p < 0.001; Table 4). These differences were again driven by *Cx. quinquefasciatus* and *Ae. aegypti*, which accounted for 23% and 8% of the differences between areas (Table 4). The turnover index was slightly higher in the gentrifying neighborhoods (0.45) than in non-gentrifying neighborhoods (0.42).

**Table 4.** Results for community composition analyses: perMANOVA, SIMPER, and turnover index at the neighborhood level. Significant *p* values have been bolded.

| perMANOVA |  |  |  |  |  |  |  |
| --- | --- | --- | --- | --- | --- | --- | --- |
|  | df | SumOfSqs | R<sup>2</sup> | F | p |  |  |
| Model | 1 | 0.63 | 0.07 | 4.71 | <0.001 |  |  |
| Residual | 63 | 8.48 | 0.93 |  |  |  |  |
| SIMPER |  |  |  |  |  |  |  |
| Species | average | sd | ratio | ava | avb | cumsum | p |
| *Cx. quinquefasciatus* | 0.23 | 0.16 | 1.38 | 760 | 536 | 0.46 | 0.11 |
| *Ae. aegypti* | 0.08 | 0.08 | 1.12 | 188 | 249 | 0.63 | **0.01** |
| *Wy. vanduzeei* | 0.06 | 0.11 | 0.55 | 1 | 126 | 0.76 | **0.002** |
| *Ae. tortilis* | 0.03 | 0.08 | 0.43 | 1 | 346 | 0.83 | **0.04** |
| *Cx. nigripalpus* | 0.02 | 0.12 | 0.24 | 1 | 1673 | 0.89 | 0.71 |
| *Ma. dyari* | 0.01 | 0.06 | 0.25 | 33 | 0 | 0.92 | 0.42 |
| **Turnover Index** |  |  |  |  |  |  |  |
| Gentrifying | 0.45 |  |  |  |  |  |  |
| Non-gentrifying | 0.42 |  |  |  |  |  |  |

## Discussion

Gentrification is occurring in cities across the United States and negatively impacts marginalized and vulnerable communities. Understanding how gentrification affects mosquito diversity in urban areas can improve our understanding of potential health risks, including the spread of mosquito-borne diseases. Our results show that mosquito alpha and beta diversity in Miami-Dade County, Florida, are greatly influenced by gentrification. Mosquito richness was lower at both county and neighborhood levels in gentrifying areas, with *Cx. quinquefasciatus* and *Ae. aegypti* driving approximately 30% of the variation in community composition. While many mosquito species disappear in the process of gentrification, *Cx. quinquefasciatus* and *Ae. aegypti* benefit from anthropogenic alterations in the environment and were the most dominant species in gentrifying urban areas of Miami-Dade.

Beta diversity differences were driven by *Cx. quinquefasciatus* at the county level and *Ae. aegypti, Wy. vanduzeei*, and *Ae. tortilis* at the neighborhood level. Vector species (i.e., *Cx. quinquefasciatus* and *Ae. aegypti*) were more abundant in gentrifying areas, which had a slightly higher turnover at the neighborhood level, indicating that gentrifying neighborhood mosquito communities have less stability in beta diversity. Non-gentrifying areas had a higher abundance of less epidemiologically relevant mosquito species (e.g., *Ae. tortilis*) as well as species commonly found in natural areas (e.g., *Wy. vanduzeei*). At both the county and neighborhood scales, species turnover remained consistent, suggesting that gentrification has a low impact on mosquito community turnover in Miami-Dade County.

The link between human health and gentrification is understudied (42), though multiple studies have shown that its effects are stratified by residents’ socioeconomic status and race (42,48), often exasperating inequalities. Socioecological factors have been recognized to influence mosquito populations, especially for species that thrive in urban areas, such as *Cx. quinquefasciatus* and *Ae. aegypti* (15–19,21,49,50). Our results indicate that at the neighborhood level, increases in home value and population density were associated with significant declines in mosquito richness, while increases in Black and Hispanic populations were associated with higher mosquito richness. Similar trends were observed at the county level, albeit at a smaller scale, suggesting that finer-scale urban dynamics play a stronger role in driving mosquito community composition. These results agree with previous studies showing that socioeconomic status (51) and gentrification (25) are drivers of mosquito community composition (52). For example, a study in Puerto Rico showed that neighborhoods with lower socioeconomic status have a greater mosquito diversity due to a higher availability of conducive aquatic habitats (16). Another study in Baltimore and Washington, DC, U.S. found high *Aedes albopictus* density in lower-income neighborhoods where trash and tire containers were more abundant (18). These findings suggest that while gentrification alters mosquito community composition, it also increases the vulnerability of these neighborhoods to mosquito vector infestations as the process intensifies.

Our study aimed to understand how gentrification influenced mosquito diversity, which can impact the epidemiology of mosquito-borne diseases. A limitation of this study is that it does not classify different types of gentrification (e.g., green gentrification vs retail gentrification). This could have an impact on results as green gentrification (i.e., an increase in parks or green spaces) would most likely favor different mosquitoes than gentrification in highly urbanized areas. We used the term gentrification in the broadest sense, i.e., a previously disinvested neighborhood characterized by social and economic changes, to have a larger sample size. Future studies should characterize gentrification in more detail to gain better insight into how mosquitoes respond to anthropogenic alterations in the environment at higher spatial resolutions.

## Conclusion

Our study shows that gentrification had a significant impact on both mosquito richness and community composition at the county and neighborhood levels. Two key mosquito vector species, *Cx. quinquefasciatus* and *Ae. aegypti*, were responsible for driving these differences with potential implications for mosquito-borne disease risk in gentrifying areas. Differences in alpha diversity were associated with changes in home value, population density, and race at the county level. At the neighborhood level, these same factors, along with age, were significantly associated with mosquito species richness. While beta diversity differed based on gentrification status, the turnover index indicated similar community stability between gentrifying and non-gentrifying areas. Future studies should consider the type of gentrification and investigate whether differences in mosquito community composition in gentrifying areas influence the risk of mosquito-borne disease.

## References

1. United Nations Department of Economic and Social Affairs. 68% of the world population projected to live in urban areas by 2050, says UN. 2018. Available from: https://www.un.org/development/desa/en/news/population/2018-revision-of-world-urbanization-prospects.html

2. Chen G, Li X, Liu X, Chen Y, Liang X, Leng J, et al. Global projections of future urban land expansion under shared socioeconomic pathways. Nat Commun. 2020;11(1): 537.

3. Gao J, O’Neill BC. Mapping global urban land for the 21st century with data-driven simulations and shared socioeconomic pathways. Nat Commun. 2020;11(1): 2302.

4. Acolin A, Crowder K, Decter-Frain A, Hajat A, Hall M, Homandberg L, et al. Gentrification yields racial and ethnic disparities in exposure to contextual determinants of health. Health Aff. 2024;43(2): 172–80.

5. Hwang J, Ding L. Unequal displacement: Gentrification, racial stratification, and residential destinations in Philadelphia. Am J of Sociol. 2020;126(2): 354–406.

6. Elliott-Cooper A, Hubbard P, Lees L. Moving beyond Marcuse: Gentrification, displacement and the violence of un-homing. Prog Hum Geogr. 2020;44(3): 492–509.

7. Lim S, Chan PY, Walters S, Culp G, Huynh M, Gould LH. Impact of residential displacement on healthcare access and mental health among original residents of gentrifying neighborhoods in New York City. PLoS One. 2017;12(12): e0190139.

8. Whittle HJ, Palar K, Hufstedler LL, Seligman HK, Frongillo EA, Weiser SD. Food insecurity, chronic illness, and gentrification in the San Francisco Bay Area: An example of structural violence in United States public policy. Soc Sci Med. 2015;143: 154–61.

9. Makhlouf MHE, Motairek I, Chen Z, Nasir K, Deo S V., Rajagopalan S, et al. Neighborhood walkability and cardiovascular risk in the United States. Curr Probl Cardiol. 2023;48(3): 101533.

10. Dragan KL, Ellen IG, Glied SA. Gentrification and the health of low-income children in New York city. Health Aff. 2019;38(9): 1425–32.

11. Gibbons J, Barton MS, Reling TT. Do gentrifying neighborhoods have less community? Evidence from Philadelphia. Urban Studies. 2020;57(6): 1143–63.

12. Harish V, Colón-González FJ, Moreira FRR, Gibb R, Kraemer MUG, Davis M, et al. Human movement and environmental barriers shape the emergence of dengue. Nat Commun. 2024;15(1): 4205.

13. Messina JP, Brady OJ, Golding N, Kraemer MUG, Wint GRW, Ray SE, et al. The current and future global distribution and population at risk of dengue. Nat Microbiol. 2019;4(9): 1508–15.

14. Centers for Disease Control and Prevention. Health Effects of Gentrification. Available from: http://medbox.iiab.me/modules/en-cdc/www.cdc.gov/healthyplaces/healthtopics/gentrification.htm

15. Hill M, Wilke ABB, Fuller DO, Kummer AG, Ventura PC, Beier JC, et al. Environmental and socio-economic determinants of mosquito vector species presence and abundance in Los Angeles County, California. 2023. Research Square. doi: 10.21203/rs.3.rs-3338430/v1.

16. Scavo NA, Barrera R, Reyes-Torres LJ, Yee DA. Lower socioeconomic status neighborhoods in Puerto Rico have more diverse mosquito communities and higher Aedes aegypti abundance. Journal of Urban Ecology. 2021;7(1): juab009.

17. Whiteman A, Loaiza JR, Yee DA, Poh KC, Watkins AS, Lucas KJ, et al. Do socioeconomic factors drive Aedes mosquito vectors and their arboviral diseases? A systematic review of dengue, chikungunya, yellow fever, and Zika Virus. One Health. 2020; 100188.

18. LaDeau SL, Leisnham PT, Biehler D, Bodner D. Higher mosquito production in low-income neighborhoods of Baltimore and Washington, DC: Understanding ecological drivers and mosquito-borne disease risk in temperate cities. Int J Environ Res Public Health. 2013;10(4): 1505–26.

19. Donnelly MAP, Kluh S, Snyder RE, Barker CM. Quantifying sociodemographic heterogeneities in the distribution of Aedes aegypti among California households. PLoS Negl Trop Dis. 2020;14(7): e0008408.

20. Wilke ABB, Chase C, Vasquez C, Carvajal A, Medina J, Petrie WD, et al. Urbanization creates diverse aquatic habitats for immature mosquitoes in urban areas. Sci Rep. 2019;9(1): 15335.

21. Wilke ABB, Vasquez C, Carvajal A, Moreno M, Fuller DO, Cardenas G, et al. Urbanization favors the proliferation of Aedes aegypti and Culex quinquefasciatus in urban areas of Miami-Dade County, Florida. Sci Rep. 2021;11(1): 22989.

22. Leisnham PT, Ladeau SL, Juliano SA. Spatial and temporal habitat segregation of mosquitoes in urban Florida. PLoS One. 2014;9(3): e91655.

23. Multini LC, Marrelli MT, Beier JC, Wilke AB. Increasing complexity threatens the elimination of extra-Amazonian malaria in Brazil. Trends Parasitol. 2019;35(6): 381–3.

24. Gigi Hoi A, Gilbert B, Mideo N. Deconstructing the impact of malaria vector diversity on disease risk. American Naturalist. 2020;196(3): E61–70.

25. Fidino M, Sander HA, Lewis JS, Lehrer EW, Rivera K, Murray MH, et al. Gentrification drives patterns of alpha and beta diversity in cities. Proc Natl Acad Sci. 2024;121(17): e2318596121.

26. Ostfeld RS, Keesing F. Effects of host diversity on infectious disease. Annu Rev Ecol Evol Syst. 2012;43: 157–82.

27. Likos A, Griffin I, Bingham A, Stanek D, Fischer M, White S, et al. Local mosquito-borne transmission of Zika Virus - Miami-Dade and Broward Counties, Florida, June-August 2016. Morbidity and Mortality Weekly Report. 2016;65(38): 1032–8.

28. Centers for Disease Control and Prevention. Data and statistics on dengue in the United States. 2024. Available from: https://www.cdc.gov/dengue/areaswithrisk/in-the-us.htmlhttps://www.cdc.gov/dengue/areaswithrisk/in-the-us.html

29. Centers for Disease Control and Prevention. Dengue. Available online: https://www.cdc.gov/dengue/index.html.

30. Shankar MB, Rodríguez-Acosta RL, Sharp TM, Tomashek KM, Margolis HS, Meltzer MI. Estimating dengue under-reporting in Puerto Rico using a multiplier model. PLoS Negl Trop Dis. 2018;12(8): e0006650.

31. Seeteram NA, Ash K, Sanders BF, Schubert JE, Mach KJ. Modes of climate mobility under sea-level rise. Environ Res Lett. 2023;18(11): 114015.

32. World Health Organization. Vector-borne diseases. 2024. Available from: https://www.who.int/news-room/fact-sheets/detail/vector-borne-diseases

33. U.S. Census Bureau. QuickFacts: Miami-Dade County, Florida. 2024. Available from: https://www.census.gov/quickfacts/fact/table/miamidadecountyflorida/POP060210

34. National Centers for Environmental Information. U.S. Climate Normals Quick Access. 2025. Available from: www.ncei.noaa.gov

35. Tedesco M, Keenan JM, Hultquist C. Measuring, mapping, and anticipating climate gentrification in Florida: Miami and Tampa case studies. Cities. 2022;131: 103991.

36. Melix BL, Jackson A, Butler W, Holmes T, Uejio CK. Locating neighborhood displacement risks to climate gentrification pressures in three coastal counties in Florida. Prof Geogr. 2023;75(1): 31–43.

37. Wilke ABB, Vasquez C, Medina J, Carvajal A, Petrie W, Beier JC. Community composition and year-round abundance of vector species of mosquitoes make Miami-Dade County, Florida a receptive gateway for arbovirus entry to the United States. Sci Rep. 2019;9(1): 8732.

38. Ventura PC, Kummer AG, Wilke ABB, Chitturi J, Hill MD, Vasquez C, et al. Forecasting the relative abundance of Aedes vector populations to enhance situational awareness for mosquito control operations. PLoS Negl Trop Dis. 2024;18(11): e0012671.

39. Darsie RF Jr, Morris CD. Keys to the adult females and fourth-instar larvae of the mosquitoes of Florida (Diptera: Culicidae). 1st ed. Vol. 1. Tech Bull Florida Mosq Cont Assoc; 2000.

40. ESRI. Esri Updated Demographics. 2024. Available from: https://doc.arcgis.com/en/esri-demographics/latest/esri-demographics/updated-demographics.htm

41. U.S. Census Bureau. 2018-2022 ACS 5-year Estimates. 2023. Available from: https://www.census.gov/programs-surveys/acs/technical-documentation/table-and-geography-changes/2022/5-year.html

42. Smith GS, Breakstone H, Dean LT, Thorpe RJ. Impacts of gentrification on health in the US: a systematic review of the literature. Journal of Urban Health. 2020;97(6): 845–56.

43. Bhavsar NA, Kumar M, Richman L. Defining gentrification for epidemiologic research: A systematic review. PLoS One. 2020;15(5): e0233361.

44. Bolker BM, Brooks ME, Clark CJ, Geange SW, Poulsen JR, Stevens MHH, et al. Generalized linear mixed models: a practical guide for ecology and evolution. Trends Ecol Evol. 2009;24(3): 127–35.

45. Chaves LF. An entomologist guide to demystify pseudoreplication: Data analysis of field studies with design constraints. J Med Entomol. 2010;47(3): 291–8.

46. Anderson M, Walsh D. PERMANOVA, ANOSIM, and the Mantel Test in the face of heterogeneous dispersions: What null hypothesis are you testing? Ecol Monogr. 2013;83: 557–74.

47. Jaccard P. Nouvelles Recherches Sur La Distribution Florale. Bulletin de la Société vaudoise des Sciences Naturelles. 1908;44: 223–70.

48. Cole HVS, Mehdipanah R, Gullón P, Triguero-Mas M. Breaking down and building up: Gentrification, its drivers, and urban health inequality. Curr Environ Health Rep. 2021;8: 157–66.

49. Ferraguti M, Martínez-De La Puente J, Roiz D, Ruiz S, Soriguer R, Figuerola J. Effects of landscape anthropization on mosquito community composition and abundance. Sci Rep. 2016;6: 29002.

50. Perrin A, Glaizot O, Christe P. Worldwide impacts of landscape anthropization on mosquito abundance and diversity: A meta-analysis. Glob Chang Biol. 2022;28: 6857–71.

51. Leong M, Dunn RR, Trautwein MD. Biodiversity and socioeconomics in the city: a review of the luxury effect. Biol Lett. 2018;14: 20180082.

52. Chaves LF, Hamer GL, Walker ED, Brown WM, Ruiz MO, Kitron UD. Climatic variability and landscape heterogeneity impact urban mosquito diversity and vector abundance and infection. Ecosphere. 2011;2(6): 70.

